# Non-resolving neuroinflammation regulates axon regeneration in chronic spinal cord injury

**DOI:** 10.1101/2024.04.19.590106

**Authors:** Andrew N. Stewart, Christopher C. Bosse-Joseph, Reena Kumari, William M. Bailey, Kennedy A. Park, Victoria K. Slone, John C. Gensel

## Abstract

Chronic spinal cord injury (SCI) lesions retain increased densities of microglia and macrophages. In acute SCI, macrophages induce growth cone collapse, facilitate axon retraction away from lesion boundaries, as well as play a key role in orchestrating the growth-inhibitory glial scar. Little is known about the role of sustained inflammation in chronic SCI, or whether chronic inflammation affects repair and regeneration. We performed transcriptional analysis using the Nanostring Neuropathology panel to characterize the resolution of inflammation into chronic SCI, to characterize the chronic SCI microenvironment, as well as to identify spinal cord responses to macrophage depletion and repopulation using the CSF1R inhibitor, PLX-5622. We determined the ability for macrophage depletion and repopulation to augment axon growth into chronic lesions both with and without regenerative stimulation using neuronal-specific PTEN knockout (PTEN-KO). PTEN-KO was delivered with spinal injections of retrogradely transported adeno associated viruses (AAVrg’s). Both transcriptional analyses and immunohistochemistry revealed the ability for PLX-5622 to significantly deplete inflammation around and within chronic SCI lesions, with a return to pre-depleted inflammatory densities after treatment removal. Neuronal-specific transcripts were significantly elevated in mice after inflammatory repopulation, but no significant effects were observed with macrophage depletion alone. Axon densities significantly increased within the lesion after PLX-5622 treatment with a more consistent effect observed in mice with inflammatory repopulation. PTEN-KO did not further increase axon densities within the lesion beyond effects induced by PLX-5622. We identified that PLX-5622 increased axon densities within the lesion that are histologically identified as 5-HT^+^ and CGRP^+^, both of which are not robustly transduced by AAVrg’s. Our work identified that increased macrophage/microglia densities in the chronic SCI environment may be actively retained by homeostatic mechanisms likely affiliated with a sustained elevated expression of CSF1 and other chemokines. Finally, we identify a novel role of sustained inflammation as a prospective barrier to axon regeneration in chronic SCI.

**Highlights:**

- Macrophages and microglia repopulate the chronically injured spinal cord after depletion.
- CSF1R antagonism in chronic SCI augments the growth of specific axon types in the lesion.
- CSF1R antagonism does not augment a PTEN-knockout-induced functional recovery.

## 1.0 Introduction

Macrophage recruitment into the lesion core is a major regulator of spinal cord injury (SCI) and recovery responses (Gensel et al., 2012; Kigerl et al., 2009; Zhang et al., 2015; Zhang et al., 2016) and is integral to the formation of the growth-inhibitory scar. In acute SCI, macrophages exacerbate axon damage through direct physical interactions and also facilitate axon die-back away from the lesion boundaries through the secretion of matrix metalloproteases (MMPs) that induce growth cone collapse and axon retraction (Busch et al., 2009; Horn et al., 2008). Indirectly, macrophages restrict axon growth and regeneration by orchestrating the formation of the glial scar (Bellver-Landete et al., 2019b; Kisucká et al., 2021; Zhou et al., 2020). Collective evidence points to inflammation as a major barrier to regeneration through the lesion via both direct and indirect mechanisms.

Signaling from macrophages orchestrate many of the glial scarring components, including the recruitment of fibroblasts into the lesion (Zhu et al., 2015), and the activation of both astrocytes and microglia (Bellver-Landete et al., 2019b; Kisucká et al., 2021; Zhou et al., 2020). Each of these cells persist at increased densities around chronic lesions and contribute to the production of a growth-inhibitory extracellular matrix (ECM) (Filous et al., 2014; Inman and Steward, 2003; Jones et al., 2003). Chronic stages after SCI are hallmarked by a stabilization of functional recovery and with very little continued resolution of the lesion environment (Beck et al., 2010; Kwiecien et al., 2020).

Interventions aiming to induce axon growth and regeneration after SCI exhibit significantly less efficacy when interventions are delayed even 1-week post-injury which corresponds to a timepoint of peak macrophage infiltration into the spinal cord (Beck et al., 2010; Blesch et al., 2012). Very little is known about the role of non-resolving neuroinflammation around chronic SCI lesions or the effects of chronic neuroinflammation on axon regeneration with, or without, growth-promoting interventions. The proceeding experiments sought to determine if persistent inflammation around and within SCI lesions function as an inhibitory barrier to axon growth in chronic SCI.

We identified non-resolving inflammation as hallmark to chronic lesions which is susceptible to macrophage/microglial depletion by treatment with the colony stimulating factor-1 receptor (CSF1R) antagonist, PLX-5622 (PLX). We combined PLX with concurrent regenerative stimulation using retrogradely transported AAV’s (AAVrg’s) to knockout the phosphatase and tensin homologue protein (PTEN) to facilitate axon growth. We observed a significant increase in axon densities within the lesion after chronic PLX treatment. However, the phenotype of axons which grew into the lesion were not those affected by AAVrg’s or PTEN-KO. Axons which expanded in the lesion were observed to be in part 5-HT^+^, but largely CGRP^+^ fibers. Collectively, we present novel data that supports inflammation within the chronically injured spinal cord as a regulator of regeneration of specific axon fiber types within the lesion.

## 2.0 Methods

### 2.1 Experimental design

#### 2.1.1 Defining the chronic SCI timeline and lesion environment

Transcriptional analysis using the Nanostring Neuropathology panel (XT-CSO-MNROP1-12; NanoString Technologies) was performed to determine a plateau in inflammatory resolution within the chronically injured spinal cord. We evaluated mRNA from spinal cord homogenates in either Naïve (n=6) mice, or at 1-week (n=6), 2-weeks (n=5), 9-weeks (n=3), and 12-weeks (n=3) post-injury. Prior reports have indicated that a resolution of inflammation reaches a plateau between 9- and 12-weeks post-SCI in rodents (Beck et al., 2010; Kwiecien et al., 2020) which was the basis of our chosen timepoints.

#### 2.1.2 Determining the ability for PLX to deplete macrophages/microglia in chronic SCI

As described below, our results validate a stable inflammatory environment by at least 9-weeks post-SCI. We sought to determine the ability for CSF1R antagonism to deplete macrophages/microglia within the chronically injured spinal cord and lesioned environment (Fig. 1). We treated mice with PLX-5622 (PLX; C-1521; Chemgood; 1200 ppm) via dietary feed (prepared at Envigo), for either 9-(n= 5), 14-(n=4), or 21-days (n=4) post-SCI. Further, we included both normal diet as a vehicle control (n=5), and a repopulation group that was treated for 21-days before allowing 3-weeks of macrophage/microglial repopulation by removing PLX (n=4). For this first experiment, mice were treated with PLX beginning at 11-weeks post-SCI, except for 1 vehicle, and 1-mouse treated for 9-days which started treatment at 7-weeks post-SCI and were used as pilot to explore treatment efficacy. These mice were combined in the groups above due to the successful depletion caused by PLX.

**Figure 1.**
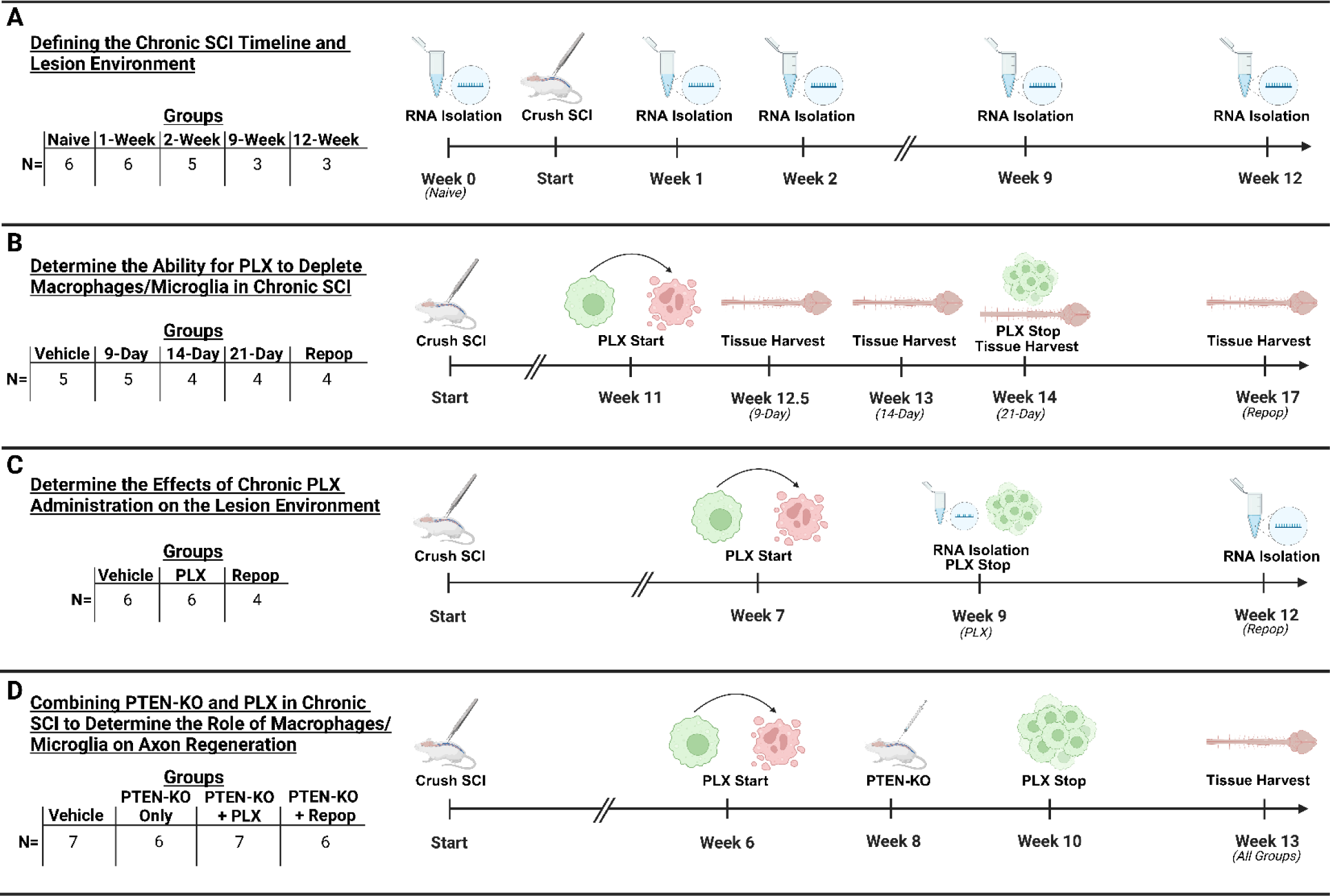
Groups, sample sizes, and timeline of experiments. In total four experiments were performed to obtain either RNA for transcriptional analysis **(A,C)**, or tissue for immunohistochemistry **(B,D)**. In our first experiment **(A)**, tissue was obtained from half the naïve mice at 9-weeks post-T9 crush SCI, and the other half at 12-weeks post-SCI for age-matched controls. In our second experiment **(B)**, tissue was obtained from one vehicle-treated mice at each tissue harvest time point, with tissue obtained from one other mouse at 9-weeks post-SCI during a pilot experiment. In our third experiment **(C)**, RNA was obtained from 3 vehicle-treated mice at week-9 and the remaining 3 at week-12 to control for time post-SCI. In our final experiment **(D)**, tissue was harvested from all mice at 13-weeks post-SCI. This figure was created with BioRender.com.

#### 2.1.3 Determine the effects of chronic PLX administration on the lesion environment

Having observed successful depletion in as little as 9-days of treatment, with better consistency observed at 14-days of treatment, we sought to determine the effects of depletion and/or repopulation on the lesioned microenvironment. We specifically hypothesized that in the absence of inflammation (i.e. while mice are sustained on PLX), tissue responses to macrophage/microglial depletion would demonstrate a reduction in inflammation with a corresponding increase in neuronal specific, or neuron-enriched transcripts. Mice were treated with either Vehicle (n=6), 14-days of PLX (n=6), or 14-days of PLX with 21-days of repopulation (n=4). For this experiment, mice began treatment at 7-weeks post-SCI, and were euthanized at either 9-weeks, or 12-weeks post-injury. For vehicle-treated mice, n=3 were euthanized at 9-weeks, and the other n=3 were euthanized at 12-weeks post-SCI to control for potential differences in the chronic injury environment. Importantly, we did not detect significant differences between 9- and 12-weeks post-SCI in any mRNA transcripts quantified after FDR correction, therefore, vehicle-treated mice from both time points were combined for analysis as a single chronic-SCI group.

#### 2.1.4 Combining PTEN-KO with PLX in chronic SCI to determine the role of macrophages/microglia on axon regeneration

To the best of our knowledge, and as was the foundation of our experimental design, axons within the chronically injured environment would likely not regenerate without a concurrent growth stimulus. Our recent experiments utilized AAVrg’s to knockout PTEN in chronic SCI, which proved to be a powerful tool to leverage growth signaling through the mTOR pathway in near-all axons damaged from SCI (Stewart et al., 2023). Importantly, however, AAVrg’s do not robustly transduce 5-HT^+^ fibers derived from the caudal raphe nucleus, nor CGRP^+^ fibers from dorsal root ganglia (DRG)(Blackmore et al., 2021; Metcalfe and Steward, 2023; Metcalfe et al., 2022; Stewart et al., 2023; Wang et al., 2018). For these reasons, we sought to determine the ability for macrophage/microglial depletion or repopulation to augment axon regeneration into the lesion when neurons are stimulated using PTEN-KO.

To get a better understanding for the role of chronic inflammation on axon regeneration, and due to compelling findings that indicated that both the pre-, post-, and repopulated-macrophage/microglial groups all display significantly different microenvironments, we advanced the following groups in this experiment: Vehicle (group name Vehicle; no PTEN-KO and no PLX; n=7), PTEN-KO/Only (group name PTEN-KO; n=6), PTEN-KO/PLX (group name PLX; sustained for 7 weeks during the experiment; n=6), and PTEN-KO/Repopulation (group name Repop; received 28 days of PLX prior to PTEN-KO, then removed from PLX treatment for 21 remaining days of the study; n=7). Mice were given PLX beginning week 6 post-SCI, AAVrg’s to deliver PTEN-KO or RFP alone at 8-weeks post-SCI and removed from PLX at 10-weeks post-SCI for the repopulation group. Mice were allowed 5 weeks post-PTEN-KO to evaluate for locomotor improvements, totaling 13 weeks post-SCI of survival (see Fig. 1 for timeline).

### 2.2 Animals and spinal cord injury modeling

All procedures were approved by the University of Kentucky Institutional Animal Care and use Committee. For experiments that did not require use of PTEN-KO, female wild-type C57BL/6J mice of age 3-months were used. For experiments that required PTEN-KO, C57BL/6J PTEN-Flox mice were used (B6.129S4-PTEN^tm1hwu^/j; strain #006440; The Jackson Laboratory) as previously described (Stewart et al., 2023). In total 108 mice were used for all experiments, with 87 mice surviving their respective experimental paradigms for data collection. Attrition was observed due to mortality during or after surgery, bladder rupture, and removal from the study for failed injections or other surgical complications such as torn dura.

Complete spinal crush injuries were performed for all mice in all experiments, using Ketamine (100.0 mg/kg) and Xylazine (10.0 mg/kg) for anesthesia. A laminectomy was performed at the T9 vertebral level, followed by a complete spinal crush produced using fine tipped forceps. The forceps were closed around the spinal cord ensuring the tips scraped the ventral side of the spinal column. Force was applied for 8 seconds before removing the forceps and closing the incision. Mice were placed on a heating pad for recovery. All mice received 1.0-mL of saline support daily for 5-days, as well as enrofloxacin at 5.0 mg/kg/day for 5-days. Buprenorphine slow-released formulation (Buprenex SR, 1.0 mg/kg) was provided once after surgery. Daily bladder expressions were provided twice each day until the end of the study.

For delivery of AAVrg’s, at 8 weeks post-SCI, mice were again anesthetized as described above and the scar tissue overlying the original laminectomy site was dissected away from the spinal cord. Mice were suspended by vertebral clips and a 10.0 µm-diameter glass pulled pipette (TIP10TW1-L; World Precision Instruments) was lowered approximately 0.5 mm into the center of the spinal cord about 1.0-mm rostral to the spinal lesion. In many instances the spinal lesions were unable to be identified, so the needle was placed as close to the T8 vertebra as possible for injections. In total, 3 mice were confirmed to have injections below the lesions and were not included in analysis, and 1 mouse presumably treated with PTEN-KO was removed for having low-to-no visible viral labeling with a concurrent surgical note questioning the success of the injection. One final mouse was removed from the study for possessing an abnormally large and visually identifiable lesion upon re-exposure, of which, spinal lesions were not typically observable after re-exposing the cord. A total of 2.0 µL of AAVrg (1.0 x 10^9^ gc/uL) carrying either Cre recombinase and dTomato (107738-AAVrg; Addgene, Watertown, MA), or mCherry alone (114472-AAVrg; Addgene, Watertown, MA) as a control, was injected into the spinal cord at a rate of 0.3 µL/min. pAAV-hSyn-Cre-P2A-dTomato was a gift from Rylan Larsen (Addgene viral prep # 107738-AAVrg; http://n2t.net/addgene:107738; RRID:Addgene_107738). pAAV-hSyn-mCherry was a gift from Karl Deisseroth (Addgene viral prep # 114472-AAVrg; http://n2t.net/addgene:114472; RRID:Addgene_114472). The needle was allowed to sit in place for 3.0-minutes to restrict backflow through the needle track, and the lesion was closed as described above. Mice received antibiotics, saline, and analgesic support as described above for this second survival surgery.

Upon completion of experiments, mice were euthanized via an overdose of Ketamine (180.0 mg/kg) and Xylazine (20.0 mg/kg) followed by cardiac perfusion with PBS and/or 4% formaldehyde prepared from paraformaldehyde powder (1558127; Sigma-Aldrich). For group assignment, animals from all experiments except the PTEN-KO and PLX combinatorial treatment were randomly assigned into groups. For our final PTEN-KO and PLX combinatorial treatment, mice were interspersed into groups at 5-weeks post-SCI prior to beginning treatment with PLX to establish a consistent mean BMS score in all groups. Small perturbations to week-5 mean scores between groups emerged from attrition due to complications arising during the re-exposure surgery for delivering PTEN-KO.

### 2.3 Transcriptional profiling of the lesioned environment

#### 2.3.1 RNA isolation

RNA was isolated for transcriptional analysis using the Nanostring Neuropathology panel from either Naïve mice, or at 1-, 2-, 9-, or 12-weeks post-SCI as well as with mice treated with PLX at 7-weeks post-SCI with/without repopulation as described above. Mice were perfused using sterile phosphate buffered saline (PBS), and approximately 5.0-mm of spinal cord surrounding the lesion was isolated and homogenized in RLT Lysis buffer containing β-mercaptoethanol (10.0 µL/mL of 14.3 µM β-mercaptoethanol; 63689; Sigma-Aldrich). Debris was pelleted and the supernatant was passed through genomic DNA eliminator columns prior to purifying on spin columns using the manufacturers protocols (RNeasy Plus; 74134; Qiagen). RNA was diluted to 13.0 ng/µL for analysis with Nanostring. All RNA was verified for degradation using Bioanalyzer prior to assessments. The Nanostring Neuropathology panel was performed at the University of Kentucky Genomics Core Laboratory following manufacturers recommendations.

#### 2.3.2 Data analysis

Data was normalized using the Nanostring N-Counter software and normalized mRNA counts were obtained. A minimal threshold count of 20 RNA copies was set as a minimal value, which represents approximately twice the value of the geometric mean of the negative controls.

Due to a copious amount of data provided by the Nanostring panel across time points and treatment conditions, we have chosen to assess and present data that is directly related to the major hypotheses. Importantly, all data will be made available through the Open Data Commons for Spinal Cord Injury (ODC-SCI). Relevant to our first objective, data was compared to evaluate for 1) establishment of a stable inflammatory environment, and 2) a deeper understanding of the chronic SCI microenvironment relative to Naïve controls. To accomplish these tasks, cell-specific transcripts of inflammation were identified and pulled from the data set and evaluated for their levels of expression at all time points. Multiple analysis of variance (MANOVA) was conducted and followed up with Dunnett’s pair-wise comparison between all other groups and the 12-week timepoint to identify significant changes relative to our last chronic SCI timepoint.

Next, both the 9- and 12-week chronic timepoints were collapsed and the data was compared to the Naïve condition to identify differentially regulated genes related to the following cellular compartments including: inflammatory-cell transcripts, neuron-enriched transcripts, secreted-protein transcripts, and extracellular matrix-related (ECM) transcripts (see Supplemental Table 1 for a list of all assessed genes). All genes used for analysis were Log2-transformed and assessed using multiple T-tests. False discovery rate (FDR) was corrected using the Benjamini and Hochberg method.

We hypothesized that depletion would elicit a response from neurons that would be consistent with regeneration or plasticity. To analyze the effects of inflammatory depletion on non-inflammatory compartments in the chronic SCI environment, we compared both mice on PLX or mice in the repopulation group to the vehicle-treated chronic SCI controls. mRNA transcripts were pulled and clustered to assess for the inflammatory, neuron-enriched, secreted, and ECM proteins described above. Each treatment was separately analyzed against the vehicle-treated chronic-SCI controls using multiple T-tests with subsequent FDR corrections. For all Nanostring analyses, only genes which possess significant discoveries are presented in figures, and all groups are presented on the same graphs.

### 2.4 Immunohistochemistry

After fixation with formaldehyde, spinal cords were isolated, acclimated to 30% sucrose solution for 1-week, frozen in blocks of 4-8 cords/block in Shandon M1 Embedding Matrix (1310; Fisher Scientific), and cut in the sagittal plane at 20.0 µm thickness at -16°C on a cryostat. Tissue sections were picked up directly on slides and labeled for immunohistochemistry. To improve antibody penetration and tissue labeling, particularly in the white matter, all slides were cleared for lipids by passing sections through graded ethanol concentrations to remove water (70%, 95%, and 100% ethanol), followed by immersing in Xylene for 5 minutes. Slides were subsequently rehydrated by passing back through the ethanol concentrations in reverse order, and re-acclimating to aqueous PBS. Slides were treated with antigen retrieval consisting of 10.0 mM sodium citrate buffer with 0.05% Tween-20 (pH 6.0) at 80°C for 5 minutes. Unspecific binding of antibodies was blocked using 5% normal goat serum (NGS) in PBS and 0.1% triton (PBS/T) for 1-hour at room temperature. All primary antibodies were incubated at room temperature overnight in PBS/T, and secondary antibodies were incubated for 1-hour at room temperature.

To assess β3-Tubulin axon growth in lesions, 1 out of every 7 sections cut in series were used for analysis. To assess all other outcomes, 1 out of every 14 sections were used for analysis. The percentage of labeled area within the lesion was used to assess all outcomes and was measured by tracing the GFAP lesion boundaries and setting an intensity threshold on the Indica Labs Halo software (Halo: Indica Labs) to measure positively labeled pixel areas. Total axon labeling was revealed using immunolabeling against β3-Tubulin (β3-Tub; 1:2,000; 5568S; Cell Signaling Tech) and was imaged using confocal microscopy. All other outcomes were imaged using conventional fluorescence microscopy on the Axioscan Z.1 (Carl Zeiss AG). Total inflammation (Iba-1; 1:4,000; 019-19741; Wako), microglia (P2ry12; 1:2,000; 94555; AnaSpec Inc.), serotonin (5-HT; 1:4,000; 20080; ImmunoStar), and calcitonin gene-related peptide (CGRP; 1:2,000; 24112; ImmunoStar) were revealed using immunolabeling. All sections were co-labeled with astrocytes (GFAP; 1:5,000; GFAP; Aves Labs) to reveal the lesion boundaries.

### 2.5 Behavioral monitoring

Mice were monitored for behavioral recovery using both the Basso Mouse Scale (BMS)(Basso et al., 2006). Two reviewers who were blinded to experimental groups assessed functional abilities as defined in the BMS scoring systems, allowing at least 4-minutes per mouse for analysis. In our prior reports we utilized the Basso, Beattie, and Bresnahan scale of locomotor recovery (BBB)(Basso et al., 1995) to assess mice with near-complete paralysis due to the increased resolution for lower-limb assessments at the lower ranges of motor abilities. We observed the ability for chronic PTEN-KO to restore weight supporting abilities in some mice, including weight support in stance, as well as several mice regaining dorsal stepping (Stewart et al., 2023). For this study, we anticipated a similar need to resolve mice transitioning from pre-to post-weight supporting abilities, and to better depict these improvements in our data. In addition to reporting the BMS scores, we used the pre-defined scoring criterion on the BMS scale to partition out a subscale that separates out plantar placement without weight support, plantar placement with weight support, and occasional to consistent dorsal stepping, all of which are clustered as a single value of 3 on the BMS. We used this Low Range Subscore to better capture the pre-to post-weight supporting transition (see Fig. 7).

### 2.6 Statistics

Statistical analysis of data from the Nanostring Neuropathology panel was specific to the objectives. First, we sought to determine the stability of inflammation within the chronic injured environment. Genes attributed to cell-specific inflammatory markers were extracted from the data set and analyzed using MANOVA with a subsequent Dunnett’s pairwise comparisons to compare between the Naïve, 1-, 2-, 9-week timepoints, to the 12-week timepoint (Performed in SPSS V28.0.0; all other analyses were performed in Prism Graphpad V10.1.1). We sought to determine the extent to which inflammatory resolution plateaus between the 9- and 12- week timepoints. Next, we wanted to identify significant differentially regulated genes. Individual T-tests were performed between the combined chronic SCI group (9- and 12-week groups) and naïve controls using the Benjamini and Hochberg method for FDR. All genes deemed significant discoveries were binned into the following categories for presentation; 1) Inflammatory-cell transcripts, 2) neuron-enriched transcripts, 3) cell-secreted transcripts, and 4) ECM transcripts (Supplemental Table 1). The same analytical strategy was applied to compare chronic SCI to PLX, and chronic SCI to mice with inflammatory repopulation.

For IHC experiments, when homogeneity of variance demonstrated significant differences between groups using the Brown-Forsythe test, Welch’s one-way ANOVA was used with Dunnett’s T3 used for pairwise comparisons. Otherwise, one-way ANOVA was used with Dunnett’s pairwise comparisons used for post-hoc testing. In our combined PTEN-KO and PLX experiments, a second analysis was performed to evaluate the main effects of PLX on axon regeneration by collapsing groups into No-PLX (vehicle and PTEN-KO mice) and PLX-treated mice (PLX and Repop groups). Combined data was assessed using Welch’s T-Test. For behavioral assessments, two-way ANOVA with repeated measures was used to compare between groups, followed by use of Fisher’s LSD for pairwise comparisons. Further, to refine our analysis of behavioral data, we assessed the max performance from all recorded time points pre-PTEN-KO against the max recorded performance post-PTEN-KO using a two-way ANOVA with post-hoc assessment using Fisher’s LSD to compare within-group and to vehicle-treated conditions. Finally, as our major conclusion supported a main effect of PTEN-KO with no effect of depletion or repopulation, we collapsed all PTEN-KO-treated groups for a refined comparison against vehicle-treated mice using two-way ANOVA with Fisher’s LSD as a post-hoc assessment.

## 3.0 Results

### 3.1 Chronic SCI lesions are hallmarked by a sustained elevation of inflammation with a persistent differential regulation in the secretome and extracellular matrix

To determine the role of chronic intraspinal macrophage activation on axon growth and regeneration, we first sought to identify a chronic timepoint in our model with non-resolving, stable, and persistent inflammation. Specifically, we utilized the Nanostring Neuropathology panel to quantify mRNA transcripts related to inflammation and neuropathology in tissue isolated from uninjured (Naïve) spinal cords or spinal cords at 1, 2, 9, and 12 weeks after severe thoracic T9 crush SCI (Fig. 1A). We defined chronicity as no significant change of inflammation between two identified timepoints.

Similar to previous reports (Beck et al., 2010; Kwiecien et al., 2020), we observed that inflammatory markers peaked between 1-2 weeks post-SCI, with a decrease to stable expression observed between 9- and 12-weeks post-SCI (Fig. 2). Only 1 out of 23 of our assessed inflammatory markers, TMEM119, changed between the 9- and 12-week timepoints (*p* = 0.039), and expression increased rather than decreased (Fig. 2). Moving forward, we collapsed the 9- and 12-week timepoints into a single chronic-SCI group for comparisons.

**Figure 2.**
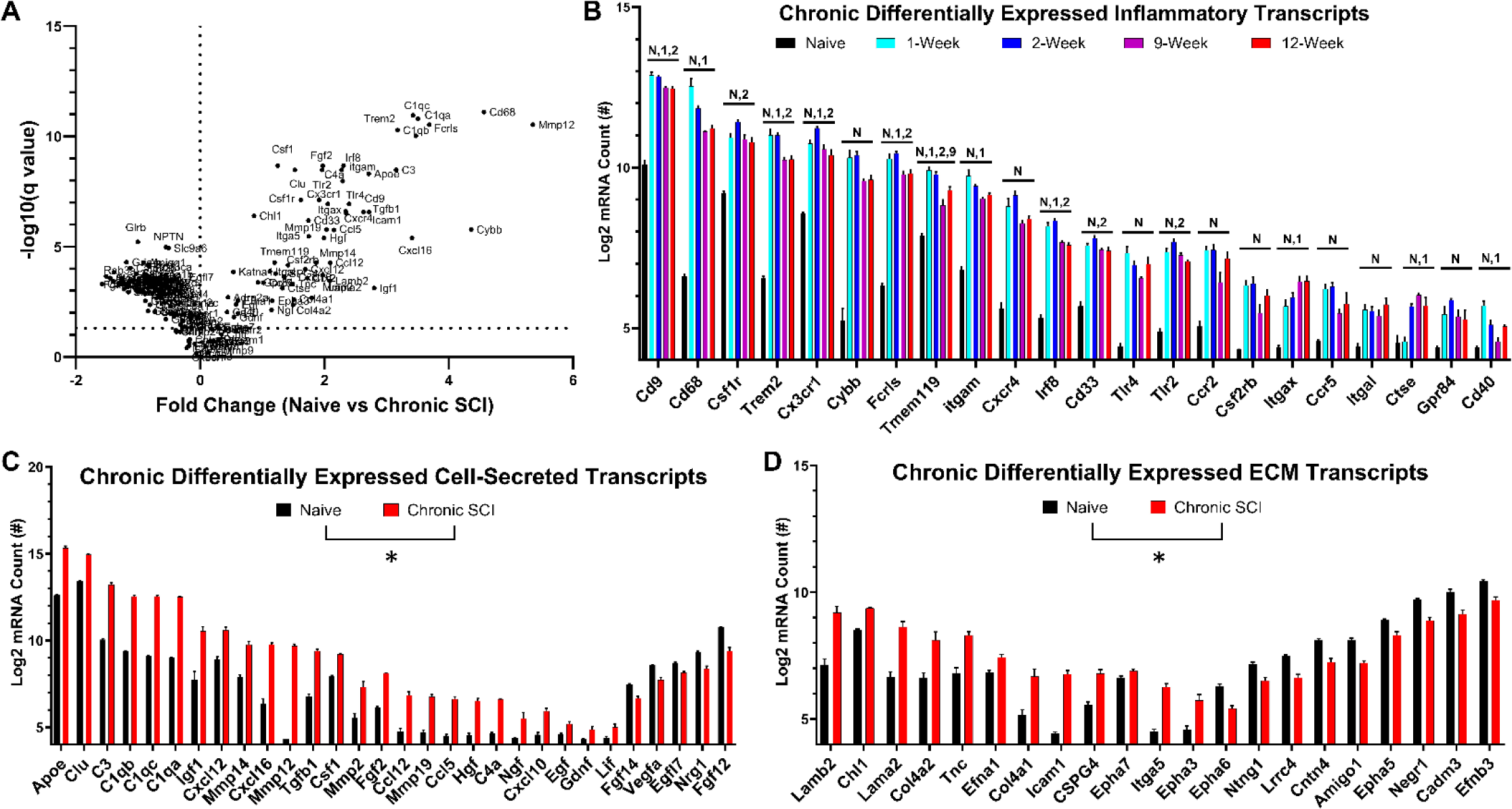
Non-resolving inflammation is hallmark to chronic SCI and retains a lesion microenvironment with differentially regulated cell-secreted and extracellular matrix transcripts. Transcriptional analysis **(A)** of the lesioned microenvironment reveals peak inflammatory profiles between 1- and 2-weeks post-SCI with incomplete resolution towards uninjured levels that plateaus between 9- and 12-weeks post-injury **(B)**. Comparing naïve to chronic-SCI lesions displays a microenvironment with persist differential regulation of the extracellular compartment including cell-secreted **(C)** and extracellular matrix-related transcripts **(D)**. Transcripts displaying a persistent upregulation in chronic SCI tend to be inflammatory-related, while genes downregulated in chronic SCI tend to be transcripts which are enriched in neurons, with diverse effects on cell-secreted and extracellular matrix transcripts **(A)**. Error bars = S.E.M. *p* < 0.05 for 12-weeks vs: N = Naïve, 1 = 1-week, 2 = 2-weeks, and 9 = 9-weeks post-SCI using MANOVA and Dunnett’s pairwise comparisons **(B)**. \**p* < 0.05 after FDR correction for all genes displayed **(C,D)**.

Of the 162 selected mRNA transcripts examined between naive in chronic SCI mice, 140 transcripts were significantly different after FDR correction. Specifically, 23/23 inflammatory, 65/66 neuron-enriched, 29/37 secreted, and 21/34 ECM mRNA transcripts were differentially regulated between naïve and chronic SCI conditions. Collectively, transcriptional analysis of tissue around chronic SCI lesions supports a persistent change in the microenvironment that may play both a role in affecting local neuronal physiology as well as affecting the potential for axon growth and repair (Supplemental Table 1).

### 3.2 CSF1R antagonism with PLX-5622 in chronic SCI depletes macrophages and microglia both within and surrounding the lesion

While we confirmed that persistent inflammation remains a hallmark of chronic SCI and stabilizes between 2 and 3 months after SCI, the influence of sustained inflammation on ongoing functions and/or tissue repair remains to be fully elucidated. To determine the role of chronic intraspinal inflammation on repair processes, we first depleted microglial and macrophages through CSF1R antagonism. Treatment with the CSF1R antagonist, PLX-5622 (PLX), was initiated at 11-weeks post-injury and maintained until spinal cord tissue was harvested either 9-, 14-, or 21-days later (Fig. 1). One additional group was allowed to recover for 21-days after removal of PLX before tissue isolation at 17 weeks post-SCI.

Histological evaluation of microglia (P2ry12) and pan macrophage (Iba-1) markers revealed significant reductions in microglia/macrophages with PLX treatment that was evident by 9 days post-treatment, with stable depletion by 14 days, and no further decrease in macrophage/microglia markers between 14- and 21-days post-treatment (Fig. 3). Of important note, not all Iba-1+ cells were depleted from within the lesion. Larger and more round-in-appearance macrophages within the lesion did not appear to deplete with PLX treatment (Fig. 3F). When PLX treatment was discontinued for three weeks, macrophages and microglia repopulated the lesion site to the same extent as vehicle-treated controls (Fig. 3A-B). For statistical comparison, we collapsed the PLX-treated groups to compare the effects of PLX to vehicle-treated mice and mice allowed for inflammatory repopulation. PLX exerted a significant depletion effect (F(_2,19_)=10.87, *p* = 0.0007) between vehicle-treated (*p* = 0.024) and repopulation-mice (*p* = 0.0007)(Fig. 3.). These effects were independently verified in a second cohort of animals from mice described in section 3.5 below (Fig. 3D). Specifically, we observed a depletion effect (F_(2,23)_=5.58, *p* = 0.01) between mice treated with PLX and those that did not receive PLX (combined vehicle-controls and PTEN-KO-treated mice)(*p* = 0.007) as well as between mice allowed for 21-days of inflammatory repopulation (*p* = 0.022)(Fig. 3D).

**Figure 3.**
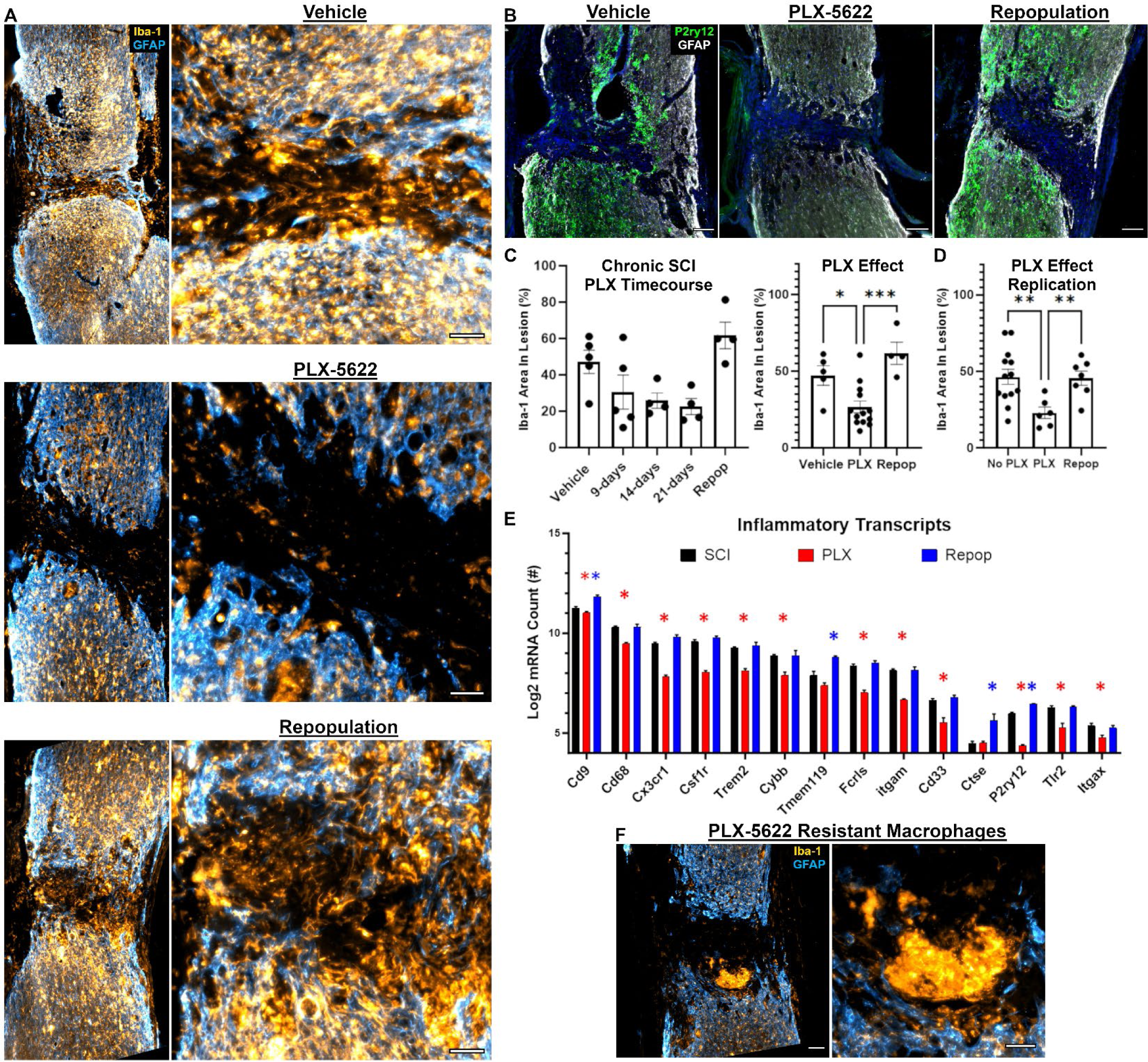
Macrophages and microglia repopulate the lesion environment back to pre-depleted levels upon removal of PLX-5622 treatment when delivered in chronic SCI. CSF1R antagonism with PLX-5622 delivered at 11-weeks **(C)**, or 6-weeks **(D)**, post-SCI depletes Iba-1^+^ macrophages and microglia from within the lesion boundaries **(A,C)**. Macrophages and microglia repopulate the lesioned environment within 21-days of removing PLX-5622 **(A,C,D)**. Microglial specific marker, P2ry12 continues to display a peri-lesional expression and low-to-no expression within the lesion core in chronic SCI **(B)**. Repopulating macrophages and microglia that re-infiltrate the lesion do not express microglial specific P2ry12 **(B)**. Transcriptional analysis of the lesioned environment demonstrates consistent findings with immunohistochemistry **(E)**. Specifically, PLX-5622 depletes inflammation across multiple inflammatory-related markers, but removal of treatment results in a restoration back to, or above, pre-depleted levels **(E)**. While PLX-5622 does significantly deplete inflammation within the lesion, some Iba-1^+^ cells display resistance to PLX-5622 and are retained during treatment **(F)**. Effects of PLX-5622 on inflammatory depletion and repopulation within the lesion were replicated in a second cohort of mice from experiments described in section 3.5 (No PLX group combined both vehicle-treated and PTEN-KO only mice) **(D)**. Scale bars = 100 µm **(B,F left)**, 50 µm **(A,F right)**. Error bars = S.E.M. \**p* < 0.05, \*\**p* < 0.01, \*\*\**p* < 0.001 **(C,D,E)**. \**p* < 0.05 after FDR correction **(E)**.

As mentioned above, upon removal of PLX, macrophages and microglia repopulated back to pre-depleted levels by 21 days after treatment removal. This was unexpected so we labeled tissue sections for the microglial protein, P2ry12, to determine if inflammation within the lesion was microglia in origin. As previously reported, macrophages within the lesion prior to depletion largely do not express P2ry12 (Stewart et al., 2021). In our study, Iba-1^+^ cells that repopulated the lesion similarly do not express P2ry12, while repopulation at the lesion penumbra do express P2ry12 (Fig. 3B). While our data do not support the repopulation of microglia into the lesioned environment, we similarly cannot conclude that repopulating macrophages are not derived from microglia in origin. Further work should identify the source of repopulating inflammation in the different compartments within the injured spinal cord (e.g. within the lesion vs penumbra).

To better characterize the inflammatory depletion and repopulation observed by histology, we replicated the depletion experiments and focused on isolating tissue 9-12 weeks post-injury to encompass stable non-resolving intraspinal inflammation (Fig. 2). As outlined in Figure 1, we treated SCI animals with PLX for two weeks, starting treatment at 7-weeks post-injury. Spinal cord homogenates encompassing the lesion were then isolated during depletion (9 weeks post-injury) or after repopulation (12 weeks post-injury) and analyzed using the Nanostring Neuropathology panel for transcriptional analysis of mRNA. A control cohort received normal chow, i.e. SCI vehicle control, with half the cohort euthanized at 9-weeks and the other at 12-weeks post-injury to control for time post-SCI. Consistent with our histology, we observed a significant decrease in 14/23 markers enriched in macrophages and microglia in the depleted group relative to SCI controls (Fig. 3E). After PLX was discontinued, all 14 transcripts returned to non-treated SCI values with two markers specific for microglia, P2ry12 (q = 0.002) and TMEM119 (q = 0.03) displaying a significant elevation above untreated levels after repopulation (Fig. 3D).

Collectively, between three separate replications and across two separate outcomes, we observed the ability for PLX to deplete inflammation both within and surrounding chronic SCI lesions, however, upon removal of treatment, macrophages and microglia repopulate back into the lesion up to, or above, pre-depleted levels. Of important note, not all Iba-1^+^ cells were depleted from within the lesion. Larger and more round-in-appearance macrophages within the lesion either did not deplete with PLX treatment or were recruited post-PLX delivery to clear apoptotic cells (Fig. 3F).

### 3.3 Inflammatory repopulation elicits global changes to the tissue microenvironment

We next examined the effect of macrophage/microglia depletion and repopulation on cell-secreted, extracellular matrix, and neuron-enriched transcripts. We hypothesized that the elimination of macrophages and microglia would allow for a reparative response of neurons and other tissues around the lesion. Contrary to our hypothesis, transcriptional analysis after inflammatory depletion and repopulation did not support a change in the microenvironment during inflammatory depletion (Fig. 4A). The only significantly affected transcripts during inflammatory depletion, compared to untreated SCI controls, were reduced inflammatory markers and closely corresponding genes that are known to be highly enriched in macrophages and microglia (e.g. complement proteins; Fig. 4C). There were no significant differences with depletion on ECM transcripts relative to untreated SCI controls (Fig. 4D). In contrast, repopulation was associated with a significant upregulation of many non-inflammatory related transcripts relative to non-treated SCI controls (Fig. 4B). Specifically, we observed significant increases in transcripts of secreted molecules (7/37) associated with cell migration and growth, e.g. fibroblast growth factor (Fgfl2), ciliary neurotrophic factor (Cntf), neuroregulin (Nrg1), etc. as well as ECM-related transcripts (9/34), e.g. neural cell adhesion molecule 1 (Ncam1), junctional adhesion molecule 3 (Jam3), cell adhesion molecule 3 (Cadm3), etc. (Fig.4).

**Figure 4.**
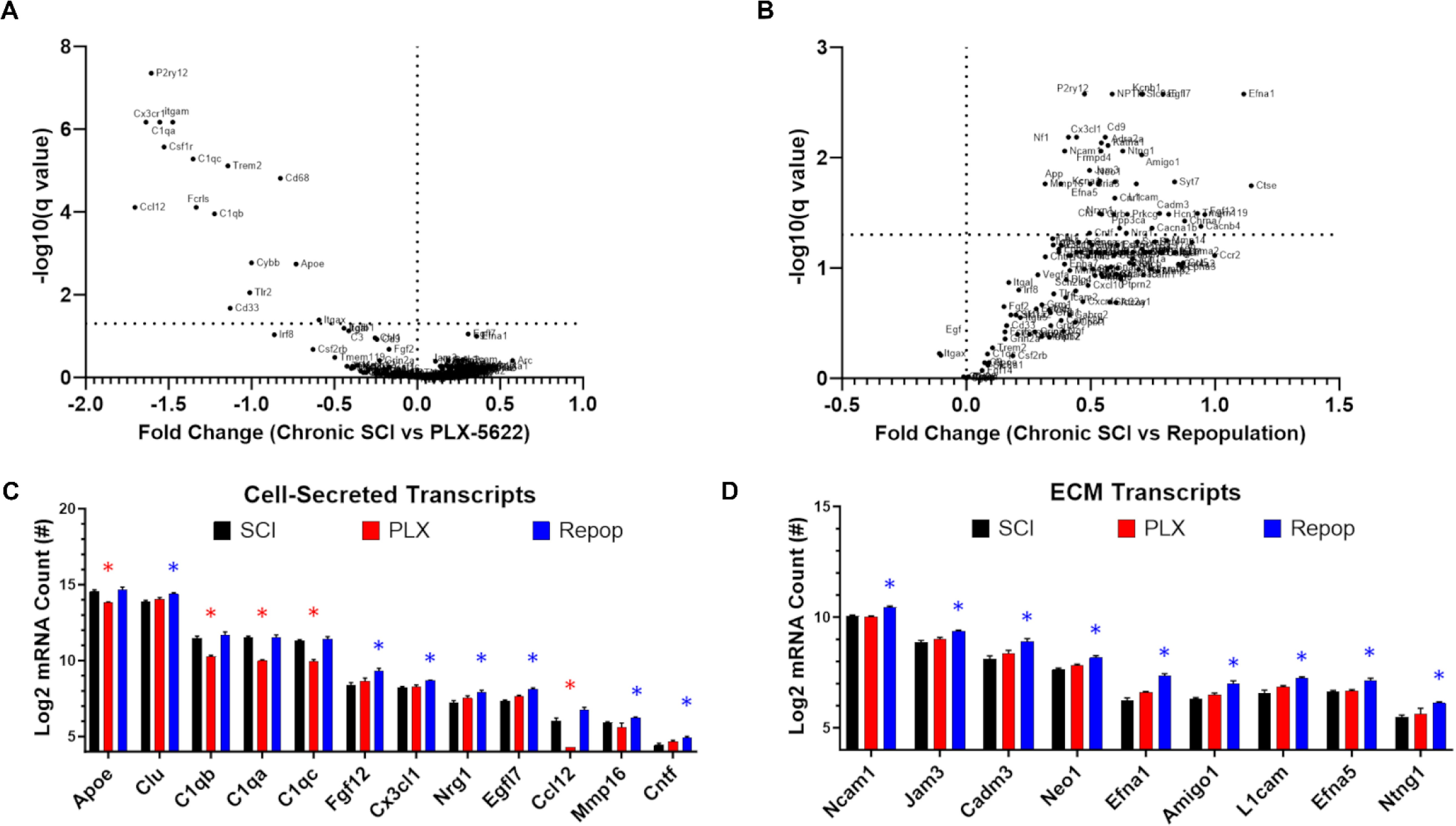
Inflammatory repopulation after PLX-5622 removal, but not depletion alone, influences the extracellular environment in chronic SCI. Delivery of PLX-5622 for 14-days starting at 7-weeks post-SCI exerts minimal effects on the extracellular environment **(A)** beyond depleting transcripts enriched in macrophages and microglia **(C,D)**. Inflammatory repopulation at 21-days after treatment removal exerted significant changes to the extracellular environment **(B)** including an upregulation of several cell-secreted and extracellular matrix transcripts **(C,D)**. Transcripts which were depleted by PLX-5622 restored back to pre-depleted values with the re-emergence of macrophages and microglia within the spinal cord **(C)**. Error bars = S.E.M. \**p* < 0.05 after FDR correction **(C,D)**.

Given the well-established interplay among inflammatory cells, secreted growth factors, and ECM proteins with neuron survival and axon growth, we next examined the effects of inflammatory depletion/repopulation on neuron-enriched transcripts. Depletion alone did not significantly alter neuron-enriched transcripts relative to non-treated SCI controls (0/66; Fig. 5A). In contrast, several neuronal transcripts were significantly increased after inflammatory repopulation relative to non-treated SCI controls (20/66). Collectively, transcriptional analysis identified that the chronic SCI environment differs depending on inflammatory depletion and/or repopulation with each treatment condition presenting a unique environment that confers a potential influence over axon growth and regeneration. Maintaining mice on PLX depletes macrophages and microglia and eliminates their contributions to the microenvironment, while repopulation of inflammation elicits a significant transcriptional response from other cells, including neurons, as well as affects several ECM-related transcripts known to play a role in axon guidance (Fig. 4).

**Figure 5.**
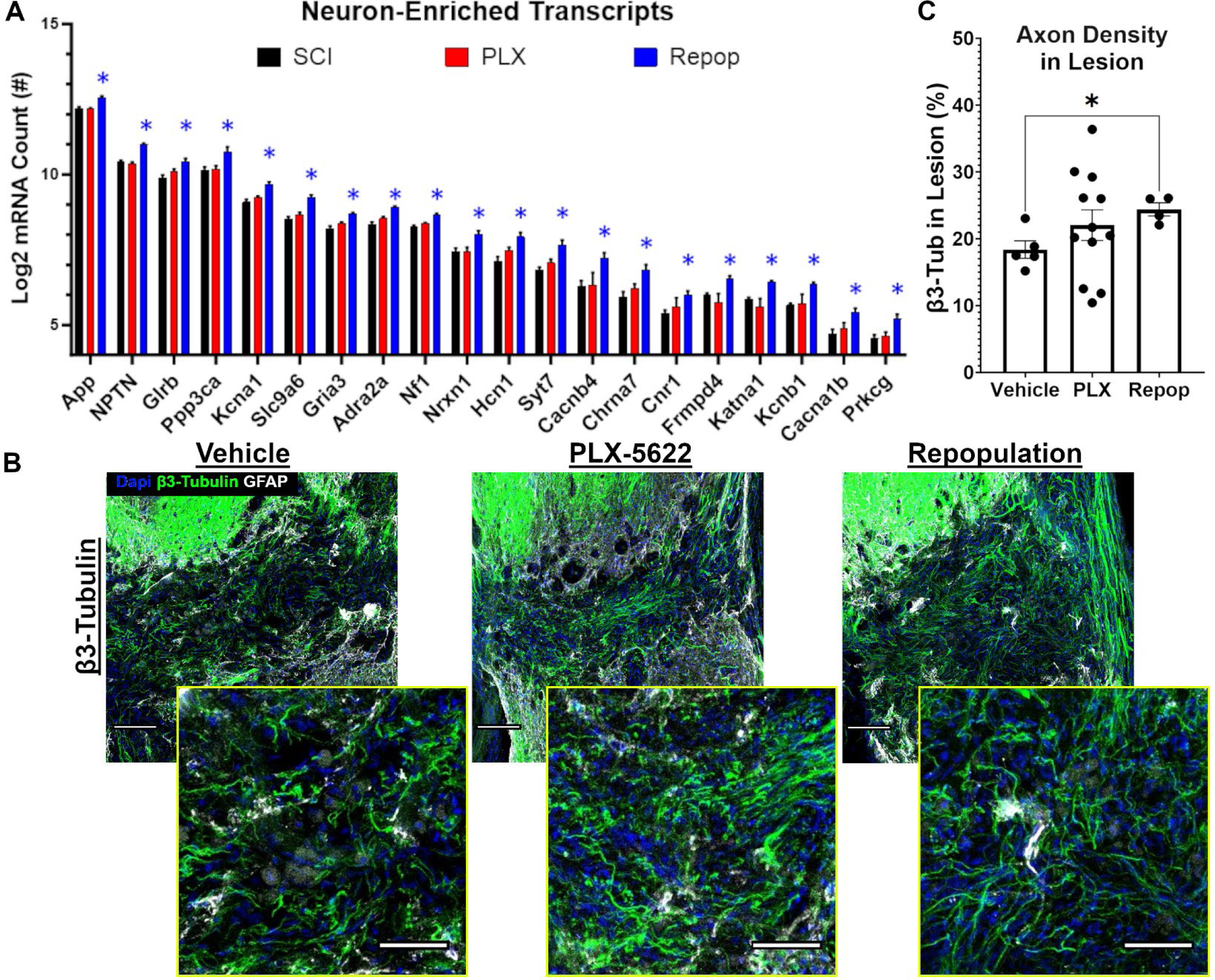
Inflammatory repopulation after PLX-5622 removal upregulates neuron-enriched transcripts within the injured environment and increases axon densities within the lesion. Lesions of mice treated with PLX-5622 and allowed for inflammatory repopulation displayed a significant upregulation of neuron-enriched transcripts **(A)**. Correspondingly, immunohistochemical analysis displayed a significant increase of axon densities within the lesions of mice allowed for inflammatory repopulation, with trends towards an increase with depletion alone **(B,C)**. Importantly, mice treated with PLX-5622 were allowed to survive for 9-21 days post-treatment initiation, while repopulated lesions were assessed 42-days post-treatment initiation and 21-days post-treatment removal. For immunohistochemical analyses, 1 mouse was euthanized at each timepoint to control for total time post-SCI. For transcriptional analyses, 3-mice were euthanized along-side mice which received 14-days of PLX-5622, and 3-mice were euthanized alongside mice that were allowed 21 of repopulation after treatment removal. Scale bars = 100 µm for less magnified images and 50 µm for zoomed images. Error bars = S.E.M. \**p* < 0.05 after FDR correction **(A)**. \**p* < 0.05 **(C)**.

### 3.4 Repopulation of macrophages elicits a neuronal response that augments axon densities in chronic SCI lesions

Having observed that macrophage depletion and repopulation significantly influence the chronic SCI environment, we sought to determine if either condition alone would augment axon growth into the lesion using histological analyses. Specifically, we labeled for the pan-neuron marker, β3-Tubulin, and assessed axon densities within the lesion. We observed a significant increase in axon densities (W(_2,10.74_) = 6.372, *p* = 0.015) in chronic SCI lesions in inflammatory repopulated mice (*p* = 0.015). While we did not detect a significant increase with depletion alone (*p* = 0.33), the range of axon densities greatly exceeded even the lesions with inflammatory repopulation (Fig. 5B,C), indicating some prospective influence of axons with depletion alone. It remains important to interpret our results with consideration that the sections with PLX-depleted lesions were obtained between 9- and 21-days post-depletion, while mice with inflammatory-repopulated lesions were obtained 42-days post-depletion. It is possible that time-post initial depletion significantly influenced axon growth into lesion which was controlled for in our next experiments.

### 3.5 Macrophage/microglia depletion and repopulation exerts fiber-type specific axon regeneration into the lesion which is independent from PTEN-KO

Our primary goal was to determine the role of sustained inflammation on axon regeneration in chronically injured spinal cord. Endogenous regeneration and repair are limited after SCI. We therefore employed a combinatorial treatment using both PTEN-KO, a well-established pro-regenerative treatment to drive axon regeneration (Danilov and Steward, 2015; Du et al., 2015; Geoffroy et al., 2015; Liu et al., 2010), and PLX, in chronic SCI. As illustrated in Figure 1D, we initiated PLX treatment 6-weeks after SCI and PTEN-KO 8 weeks after injury. Tissue was then harvested at a timepoint indicative of chronic, sustained intraspinal inflammation (13 weeks post-injury). We included both a group with mice sustained on PLX for 7-weeks throughout PTEN-KO and as well as a group treated with PLX for only 4-weeks which allowed for 21-days of inflammatory repopulation in the presence of PTEN-KO. We used spinal injections of retrogradely transported AAVs (AAVrg’s) as previously described (Metcalfe and Steward, 2023; Stewart et al., 2023) to knockout PTEN as a growth-promoting stimulus that targets most spinal-projecting neurons descending from the brain and brainstem. Control groups included animals without PLX or PTEN-KO (aka vehicle SCI) and animals treated for PTEN-KO without macrophage depletion (PTEN-KO alone).

To first evaluate overall axon growth, regardless of phenotype, we labeled for β3-Tubulin to visualize all axons. We replicated the main effect of PLX on axon growth within the lesion when collapsing groups into those that either did or did not receive PLX (*p* = 0.002)(Fig. 6B). When assessing all 4 groups individually, we observed a significant main effect of treatment (W(_3,10.97_) = 4.544; *p* = 0.026) with the greatest difference between PTEN-KO alone vs. PTEN-KO with PLX administration with or without repopulation (Fig. 6C).

**Figure 6.**
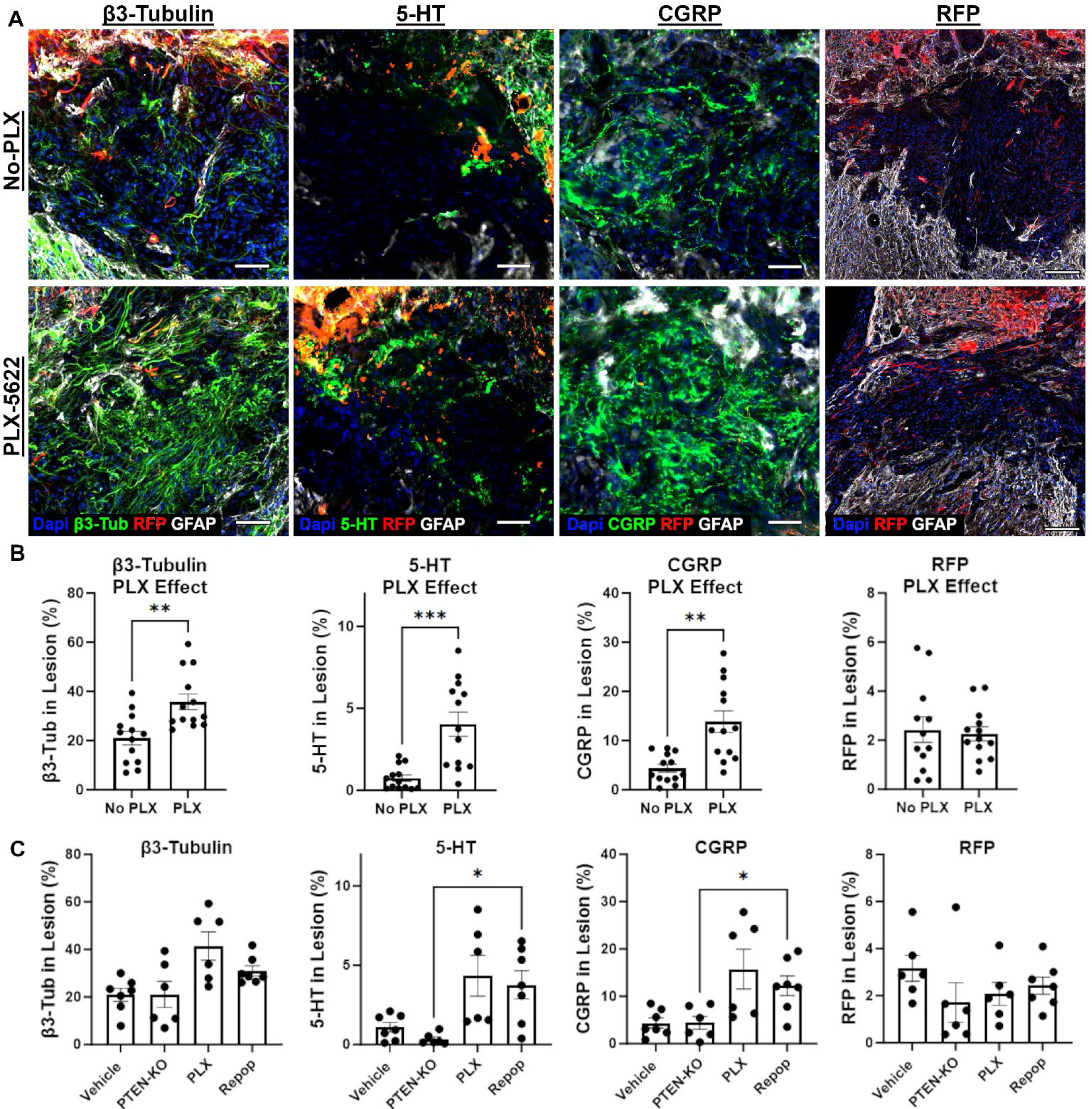
5-HT^+^ and CGRP^+^ axons expand within PLX-5622 treated chronic SCI lesions, but PTEN-KO with AAVrg’s does not further augment axon growth into the lesion. Mice were treated for 4-(Repop group; 4-weeks of treatment 3-weeks of treatment recovery) or 7-weeks (PLX group) with PLX-5622 or a regular diet as a vehicle control, beginning at 6-weeks post-SCI. Mice were subsequently subject to PTEN-KO or a vector control using AAVrg’s to deliver cre-recombinase and RFP, or RFP alone at 8-weeks post-SCI (2-weeks post-PLX-5622 initiation; Fig. 1D). Total axon growth was expanded in the lesions of mice treated with PLX-5622 regardless of allowing for inflammatory repopulation **(A,B)**. No significant increase in RFP^+^ axon growth was observed in the lesions of any PTEN-KO group relative to vehicle controls, indicating that PLX-5622 does not augment regeneration of axons susceptible to AAVrg’s **(A,B,C)**. Both 5-HT^+^ and CGRP^+^ axons were observed to significantly grow within chronic SCI lesions in response to PLX-treatment, independent of inflammatory repopulation **(A,B)**. A significant increase was observed between mice allowed for repopulation and PTEN-KO controls at the level of individual comparisons **(C)**. Scale bars = 50 µm. Error bars = S.E.M. \**p* < 0.05, \*\**p* < 0.01, \*\*\**p* < 0.001.

Observing an increase in axons within the lesions of mice treated with PLX replicated our prior experiment, however, here we demonstrate that axon growth is independent of sustained depletion or repopulation when total time post-depletion was controlled (Fig. 6B). Next, we investigated axon growth of specific populations including serotonin expression axons (5-HT) and calcitonin gene-related peptide (CGRP) positive axons which are not efficiently transduced with AAVrg’s. We detected a significant main effect of PLX treatment with increased axon growth of both 5-HT (*p* = 0.007) and CGRP (*p* = 0.001) when collapsing groups into those that did or did not receive PLX (Fig. 6B). By comparing PTEN-KO-treated mice to all other groups, we identified a significant increase in 5-HT^+^ (W(_3.0,10.48_) = 8.01, *p* = 0.0046) axon densities within the lesions of mice allowed for inflammatory repopulation (*p* = 0.023), but not in mice with sustained inflammatory depletion (*p* = 0.07)(Fig. 6C). Similar observations were made when evaluating fibers histologically identified as CGRP^+^ (W(_3.0,11.23_) = 5.34, *p* = 0.015), with a significant increase in axon densities observed between PTEN-KO and inflammatory-repopulation conditions (*p* = 0.031), but not with depletion alone (*p* = 0.1) (Fig. 6C). Interestingly, neither depletion (*p* = 0.97) nor repopulation (*p* = 0.81) affected RFP^+^ axon growth into the chronic lesioned environment beyond mice treated with PTEN-KO (Fig. 6), indicating that PLX may augment growth of specific axon sub-types that possess pre-existing abilities for limited growth into the lesion (Inman and Steward, 2003).

Collectively, we have reproduced an effect of PLX on axon growth into chronic SCI lesions with both sustained and repopulated inflammatory conditions, however, macrophage repopulation elicits a more consistent axon growth response. While we did not observe a significant augmentation of retrogradely labelled-RFP^+^ fibers, this does not preclude the possibility that axons transduced by AAVrg’s could have grown around the lesion and may not be capable of intra-lesion growth, despite inflammatory depletion. Here we identify at least two histologically identifiable axons with some potential for growth into the lesion that respond to macrophage/microglia depletion with significant growth within the lesion.

### 3.6 Inflammatory repopulation with PTEN-KO improves locomotor functions

We monitored for potential synergistic effects on locomotor recovery derived from combining PTEN-KO and PLX with and without repopulation. As a main effect over the duration of the study, we did not detect significant between-group differences (F(_3,22_) = 0.893, *p* = 0.46), nor a significant time-by-group interaction (F(_24,176_) = 1.124, *p* = 0.32). However, at the level of pair-wise comparisons, we did observe significant differences between the inflammatory-repopulation group and vehicle-controls at 3-weeks post-treatment (*p* = 0.023) using the BMS as a main outcome (Fig. 7A). Not observing a significant improvement as a main effect on the BMS was surprising considering we observed several mice in all groups receiving PTEN-KO regain weight supporting abilities at some point after treatment (6/19 mice treated with PTEN-KO with or without PLX).

**Figure 7.**
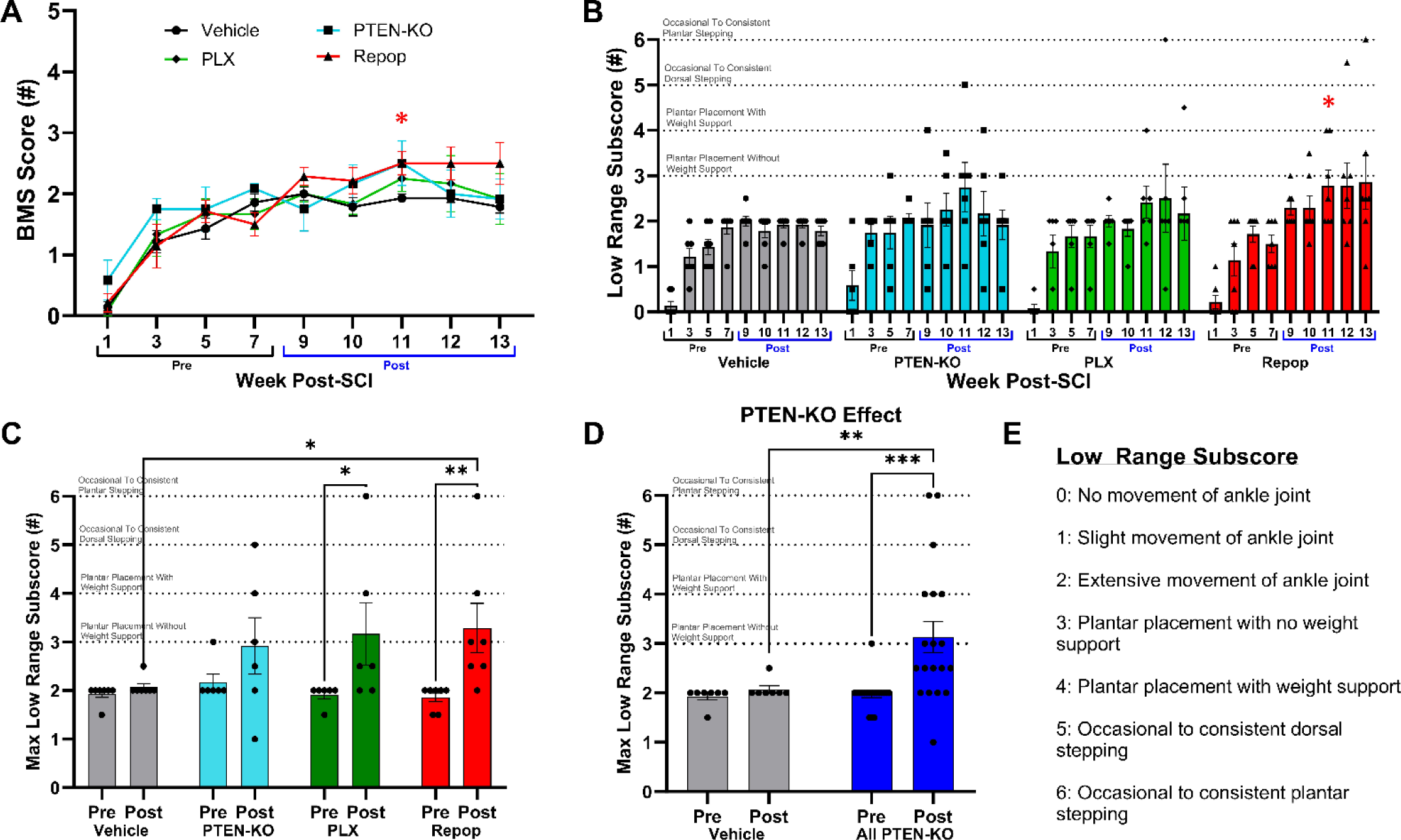
PTEN-KO with AAVrg’s restores weight support in chronic SCI but PLX-5622 does not significantly affect locomotor functions. Mice were treated for 4-(Repop group; 4-weeks of treatment 3-weeks of treatment recovery) or 7-weeks (PLX group) with PLX-5622 or a regular diet as a vehicle control, beginning at 6-weeks post-SCI. Mice were subsequently subject to PTEN-KO or a vector control using AAVrg’s to deliver cre-recombinase and RFP, or RFP alone at 8-weeks post-SCI (2-weeks post-PLX-5622 initiation; Fig. 1D). PLX-5622 treatment did not significantly affect functional recovery beyond what was observed with PTEN-KO alone. No main effects were found using two-way ANOVA for the BMS **(A)**, despite observing 6/19 mice treated with PTEN-KO regain weight support of some kind after treatment **(B)**. A Low Range Subscore was generated to better represent mice transitioning from pre-to-post weight supporting abilities using pre-defined BMS criterion **(E)**. When observing the maximal recorded performance from all pre- and post-PTEN-KO mice, within-group improvements were detected for both mice sustained on PLX and allowed for repopulation, but no differences were observed above PTEN-KO only **(C)**. The main conclusion from behavioral assessments supports the ability for chronic PTEN-KO using AAVrg’s to transiently improve locomotor functions **(D)**, but PLX-5622 does not significantly affect motor abilities in near-complete chronic crush SCI models. Error bars = S.E.M. \**p* < 0.05, \*\**p* < 0.01, \*\*\**p* < 0.001.

To gain better resolution that can capture the pre- to post-weight supporting hindlimb abilities, we generated a Low Range Subscale based on BMS scoring criterion (Fig. 7E). Prior to treatment, only one animal across all groups demonstrated hindlimb function better than extensive ankle movement (plantar placement of the hindlimb, PTEN-KO alone group). After intervention, there were no significant improvements in the vehicle-only group with only two animals (2/7, 29%) showing any increased function at any time after treatment (*p* = 0.75)(Fig. 7). Although more animals recovered function in the PTEN-KO alone group (3/6, 50%) the maximal performance before vs after treatment was not statistically significant (*p* = 0.14). In contrast, in both the macrophage-depleted and macrophage-repopulated groups, significant improvements were detected as a maximal recorded performance pre- vs post-treatment (*p* = 0.02 and *p* = 0.005, respectively) (Fig. 7C). Specifically, 5 of 6 animals in the PLX-treated group improved with treatment (83%) and 7 of 7 in the depletion and repopulation groups (100%). Relative to vehicle-control treated animals, only the repopulation group differed between vehicle-controls at 11 weeks post-treatment (*p* = 0.046), as well as in their max performance post-treatment (*p* = 0.039)(Fig. 7B). While our data implicate most of the treatment effect is likely derived from PTEN-KO (Fig. 7D), mice receiving both PTEN-KO with inflammatory repopulation displayed the largest effect that was significantly different between vehicle-treated controls.

## 4.0 Discussion

Our work presents several novel findings that implicate sustained inflammation in chronic SCI lesions as regulators of axon growth and regeneration. First, by cycling PLX to induce inflammatory depletion and repopulation, we revealed that inflammation returns back to pre-depleted levels, which are significantly elevated compared to naïve conditions. Second, 2-weeks of inflammatory depletion alone exerted minimal-to-no detectable effects on surrounding tissue other than a subsequent reduction in inflammatory-related contributions to the lesioned environment. In contrast, allowing for inflammatory repopulation elicits a significant response from non-inflammatory related cells and constituents of the extracellular environment. Third, two separate experiments of cycling PLX revealed that macrophage depletion/repopulation increases axon growth into the lesion. However, the augmentation of growth appears restricted to axon phenotypes with a pre-established capability for intra-lesion growth. Fourth, while conditions of inflammatory repopulation combined with PTEN-KO did improve motor recovery in chronic SCI, neither inflammatory depletion or repopulation significantly augmented functional improvements beyond effects of PTEN-KO alone.

A persistent increase in inflammation after SCI has been documented across most mammalian species including humans (Beck et al., 2010; Kwiecien et al., 2020; Zrzavy et al., 2021). While much research has been performed to understand the role of acute inflammation on injury progression and pathology, less has been performed to understand the role of chronic inflammation after SCI. In contrast to prior reports that have presented little evidence to support the ability for PLX to reduce inflammation within the lesions after acute-SCI (Bellver-Landete et al., 2019a; Brennan et al., 2022; Jakovčevski et al., 2021; Li et al., 2020), we observed a significant depletion effect when treatment was delayed into the stable chronic condition in our crush model. The ability for PLX to deplete macrophages/microglia from the lesioned environment depends on the reliance on CSF1R for survival. Whether or not macrophage populations within the lesion core transition to a dependence on CSF1R from acute to chronic SCI remains unknown. Our work raises support for an emerging hypothesis, specifically supporting a potential role for ongoing macrophage turnover within the chronic lesion that feeds a persistent increase of inflammatory densities.

At least two potential hypotheses exist that can explain the persistence of increased inflammation after SCI: 1) macrophages that infiltrate the lesion core become trapped within the lesion and undergo phenotypic changes throughout the progression of SCI pathology; or 2) macrophages within the lesion undergo life cycles of death and proliferation/repopulation at a rate which perpetuates sustained inflammation. The first hypothesis would be supported by an environment rich in ECM that prohibits growth and migration of tissue, as well as a well-characterized transition of macrophages towards a foam-cell phenotype that restricts cellular migration (Zhu et al., 2017), similar to macrophage dynamics in atherosclerosis (Luo et al., 2017). In our work, after significantly depleting inflammatory populations from the lesion core, we observed a repopulation of inflammation back to elevated and pre-depleted levels. While repopulation of tissue-resident macrophages has already been described after removal of PLX in other nervous system and systemic tissues (Henry et al., 2020; Lei et al., 2020), we speculated that the extent of repopulation would approach levels closer to naïve conditions. Observing a repopulation back to pre-depleted levels, which are significantly elevated over naïve conditions, supports a role of the chronic SCI environment in sustaining elevated levels macrophages/microglia around the lesion as a new homeostasis. Such implications support the hypothesis that macrophages/microglia may undergo ongoing turnover within the lesioned environment, and that other cells surrounding the lesion actively support a chronic increase in macrophage/microglia presence.

Transcriptional analysis of the chronic SCI environment revealed a significant and persistent elevation of CSF1 (Fig. 2C), which was not significantly affected by macrophage depletion. A chronic elevation of CSF1 could explain an increased support for macrophage/microglia populations within and surrounding the lesion. We originally hypothesized that a persistent increase in inflammation was supporting chronic activation of the glial scar and a persistent expression of inhibitory ECM such as NG2/CSPG4. However, our data point to the contrary, specifically supporting that non-inflammatory components of the lesioned environment sustain increased inflammation. Transcriptional analysis after inflammatory depletion did not reveal a significant effect on any cellular marker that could be attributed to non-inflammatory components of the lesioned environment. Instead, we observed a decrease in macrophage/microglia-specific markers, and their known contributions to the secretome such as complement proteins. Importantly, one recently published paper has suggested that delivery and sustaining CSF1R antagonists in chronic SCI for 10 weeks can resolve astrogliosis (Xia et al., 2022). While our data does not replicate such a conclusion, our transcriptional analysis experiments only sustained mice for 2 weeks, which may provide insufficient time for the glial scar to resolve. Overall, our findings help frame our understanding of chronic SCI pathology by revealing a prospective directionality of the relationship between sustained inflammation and the non-inflammatory components of the glial scar.

Observing that inflammatory repopulation elicited a remarkable upregulation of genes that are not affected by depletion alone implicates the potential for pro-inflammatory events in chronic SCI to regulate neural responses that may include repair and regeneration. Specifically, while inflammatory transcripts returned to pre-depleted levels, transcripts which are enriched in neurons, as well as secretome- and ECM-related transcripts, were upregulated after removal of PLX. The reason for a global response to repopulation remains unknown, however, it can likely be speculated that an environment filled with proliferating cells may increase the secretion of growth/trophic factors which would interact with other cells in the environment.

Of potential importance was a significant upregulation of MMP16 and a notable but insignificant upregulation of MMP14 (*p* = 0.058 after FDR correction; Supplemental Table 1) during inflammatory repopulation. While acknowledging the limitations that transcriptional assessments of MMPs does not equate to protein expression or activation, we recognized that inflammatory repopulation leading to MMP16/14 activity could have important implications. Specifically, the growth-inhibitory proteoglycan, NG2, is a known substrate for MMP14 (Nishihara et al., 2015), of which NG2 is upregulated in chronic SCI lesions and contributes to regenerative failure (Filous et al., 2014; Ughrin et al., 2003). There logically would be a need for proliferating and repopulating macrophages/microglia to digest ECM during their migration within the injured environment. If repopulating inflammation results in a digestion of inhibitory ECM, such a condition could inadvertently augment the potential for regeneration by removal of inhibitory scar components. The role of inflammatory events in the CNS during chronic stages of SCI has been previously explored for the ability to promote periods of plasticity that can be fostered for functional recovery but the underlying mechanisms remain to be elucidated (Torres-Espín et al., 2018).

While our original hypothesis speculated that the presence of inflammation acted as a barrier to regeneration, our strongest evidence supports that allowing for repopulation, rather than depletion alone may exert a more consistent effect and a more growth-permissive response to surrounding tissue. While we did reproduce an effect of PLX on axon growth into the lesion across two experimental replicates, surprisingly, we did not observe any evidence to support the ability of PTEN-KO (RFP^+^ fibers) to augment growth within the lesion boundaries. Such findings suggest that while axon growth was enhanced by PLX treatment, growth was occurring from axon-subtypes which are not readily transduced by AAVrg’s. Importantly, because PTEN-KO was irrelevant to intralesional growth, and because both PLX-treated groups exhibited comparable effects, we collapsed groups into PLX-treated and non-treated groups to better represent the effects of treatment. We observed a significant increase in total axon densities within the lesion, which while does replicate our first experiment does not describe the axon sub-types receptive to PLX.

Prior reports, as well as our own, describe a limited ability for AAVrg’s to transduce some axon-subtypes, including 5-HT^+^ axons descending from the caudal Raphe nucleus, as well as CGRP^+^ fibers from the dorsal root ganglia (Blackmore et al., 2021; Metcalfe and Steward, 2023; Metcalfe et al., 2022; Stewart et al., 2023; Wang et al., 2018). Importantly, both axon subtypes are known to possess some potential to growth into the lesion boundaries (Inman and Steward, 2003). Our data identifies that at least 5-HT^+^ and CGRP^+^ fibers respond to PLX with growth within the lesion. While we can conclude that PLX does augment axon regeneration of specific fiber types and can further conclude that depletion/repopulation does not support regeneration of AAVrg^+^ axons into the lesion, the ability for PLX to augment PTEN-KO-affected axons to grow around the lesion and through spared tissues was not, and cannot be, determined in our experimental design.

Prior work has suggested that regenerating corticospinal tract axons after PTEN-KO follow astroglial bridges or routes of spared tissue, rather than growing into the lesion core (Du et al., 2015; Liu et al., 2010; Zukor et al., 2013). While our data can conclude that macrophages are not causal to this failure to grow through the lesion, we were not able to identify if a subsequent regenerative response occurred through small rims of spared tissue or through astroglial bridges. Our analysis focused on growth within the lesion and did not include regions of spared tissue that were sparse but occasionally apparent via continuous GFAP boundaries. Further, as described in our prior work, while AAVrg’s are emerging to be a powerful tool to target an expanded population of descending motor neurons, some limitations to the use of AAVrg’s exist (Stewart et al., 2023). Specifically, the profound ability for AAVrg’s to target most descending axons includes spared fiber tracts that are difficult to discriminate against possibly regenerated fibers. Second, we observed neuron cell bodies caudal to the lesion. It is impossible to distinguish between neurons caudal to the lesion as having spared axons ascending through the lesion and into the injection site, vs the possibility that virus diffused across the lesion and labeled some neurons. Both complications result in the identification of RFP^+^ axons caudal to the lesion that we cannot accurately distinguish as either being regenerated, spared, or ascending in nature, which challenge interpretation for concluding regeneration.

As discussed in our prior work, it is likely that the functional benefits observed after chronic PTEN-KO are elicited via spared pathways, rather than regeneration, which is supported by the speed at which recovery occurs after treatment (Stewart et al., 2023). In both our prior work, and work presented here, we observed recovery occurring in as little as a couple weeks post-treatment, which is a rate likely too fast to be attributed to regeneration. While we did replicate a PTEN-KO-induced recovery effect, neither sustained inflammatory depletion or repopulation significantly affected locomotor abilities in either direction relative to PTEN-KO alone.

Thus, while chronic neuroinflammation may play a role in regulating axon growth and regeneration, at least within our experimental conditions, we were not able to implicate sustained inflammation as a key regulator of motor functions. It should be appreciated, however, that the severity of our injury model leaves few, if any, spared axons whereby microglial depletion at distances far away from the lesion may exert effects on sprouting or plasticity. Further work should explore the effects of macrophage/microglia depletion on functional abilities on more incomplete injury models, as well as examine other functional modalities such as sensation and pain.

### 4.1 Conclusions

In addition to concluding that sustained inflammation within chronic SCI lesions negatively regulates axon regeneration of specific fiber types, we have identified that an undetermined homeostatic mechanism retains increased macrophage and microglial densities within and around the lesion. Our results reveal a novel role of the lesion in sustaining chronic inflammation and identifies non-resolving inflammation as a regulator of axon growth and regeneration.

## Abbreviations

5-HT: serotonin
AAV: adeno-associated virus
AAVrg: retrograde psuedotyped adeno associated virus
ANOVA: analysis of variance
β3-Tub: beta III tubulin
BMS: Basso Mouse Scale
CNS: central nervous system
CGRP: calcitonin gene-related peptide
CSF1R: colony stimulating factor-1 receptor
GFAP: glial fibrillary acidic protein
hSyn1: human synapsin 1 promoter
IHC: immunohistochemistry
KO: genetic knockout
mTOR: mammalian target of rapamycin
PBS: phosphate buffered saline
PLX: the CSF1R antagonist PLX-5622
PTEN: phosphatase and tensin homolog protein
RFP: red fluorescent protein
SCI: spinal cord injury

## Data Sharing

All data from this manuscript will be publicly available through the Open Data Commons for Spinal Cord Injury (www.odc-sci.org).

## Authorship Contributions

**Conceptualization:** A.S. & J.G. **Methodology:** A.S., R.K., C.B., K.P, V.S., W.B **Validation:** A.S., J.G. **Formal Analyses:** A.S., J.G. **Investigation:** A.S., J.G. **Data Curation:** A.S., J.G. **Writing:** A.S., J.G. **Project Administration:** A.S., J.G. **Resources:** J.G. **Funding Acquisition:** A.S., J.G. **Supervision:** J.G.

## Acknowledgements

Thank you to the Wings for Life Foundation for providing funding to support this project. Confocal microscopy was performed at the University of Kentucky’s Light Microscopy Core. pAAV-hSyn-Cre-P2A-dTomato was a gift from Rylan Larsen (Addgene viral prep # 107738-AAVrg; http://n2t.net/addgene:107738; RRID:Addgene_107738). pAAV-hSyn-mCherry was a gift from Karl Deisseroth (Addgene viral prep # 114472-AAVrg; http://n2t.net/addgene:114472; RRID:Addgene_114472).

